# Overexpression of RsMYB1 enhances heavy metal stress tolerance in transgenic petunia by elevating the transcript levels of stress tolerant and antioxidant genes

**DOI:** 10.1101/286849

**Authors:** Trinh Ngoc Ai, Aung Htay Naing, Byung-Wook Yun, Chang Kil Kim

## Abstract

The RsMYB1 transcription factor (TF) controls the regulation of anthocyanin in radish (*Raphanus sativus*), and its overexpression in tobacco and petunia strongly enhances anthocyanin production. However, no data exists on whether RsMYB1 is involved in the mechanism that leads to abiotic stress tolerance. Under normal conditions, transgenic petunia plants expressing RsMYB1 and WT were able to thrive by producing well-developed broad leaves and regular roots. In contrast, a reduction in plant growth was observed when they were exposed to heavy metals (CuSO_4_, ZnSO_4_, MnSO_4_, and K_2_Cr_2_O_7_). However, RsMYB1-overexpressing plants were found to be more tolerant to the stresses than the WT plants because the expressions of stress tolerant genes (*GSH* and *PCs*) and antioxidant genes (*SOD, CAT*, and *POX*) were enhanced. In addition, according to the phylogenetic analysis, RsMYB1 has a strong sequence similarity with other MYB TFs that confer different abiotic stresses. These results suggest that overexpression of RsMYB1 enhances the expression levels of metal-induced stress tolerance genes and antioxidant genes, and the resultant increase in gene expression improves heavy metal stress tolerance in petunia.

## Introduction

Heavy metals naturally occur in the earth’s crust. However, excess levels of the heavy metals due to natural or anthropogenic activities negatively affect living organisms. Over the past few decades, industrialization and the application of modern agricultural practices has increased worldwide, which has led to the contamination of cultivatable land with heavy metals released from agro-chemicals and from industrial activities (Yang et al., 2005). Generally, some heavy metals, such as Zinc (Zn), copper (Cu), and manganese (Mn), play an important role in plant physiological and biochemical processes, such as chlorophyll biosynthesis, photosynthesis, and DNA synthesis, etc. (reviewed by Singh et al., 2015). For example, Zn has roles in the maintenance of membrane integrity, auxin metabolism, and reproduction because it interacts with enzymes and transcription factors associated with the mechanisms underlying these processes (Williams and Pittman, 2010; Prasad, 2012; Ricachenevsky et al., 2013). However, the toxic effects of the heavy metals at elevated concentrations have also been well documented (Fontes and Cox, 1998, Lewis et al., 2001, Warne et al., 2008). Zinc at elevated concentrations can stop plant metabolic functions and causes growth retardation and senescence (Fontes and Cox, 1998, Warne et al., 2008). It has also been reported that high Cu concentrations cause a similar range of symptoms (Lewis et al., 2001). In addition, elevated Mn concentrations are toxic to many plant species (Rezia et al 2005; Izaguirre-Mayoral et al. 2008) and elevated Cr negatively affects cell division, and root and stem growth in many plants (Shanker et al., 2005, Zou et al., 2006, Fozia et al., 2008). Overall, these metals at elevated concentrations lowered biomass accumulation and crop productivity by inhibiting several plant mechanisms. Theoretically, the presence of excess heavy metals limits CO_2_ fixation and reduces the photosynthetic electron transport chains in chloroplasts and mitochondria, which leads to the generation of reactive oxygen species (ROS) that damage plant cells and growth, which leads to reduced crop yields (Davidson and Schiestl, 2001, Mittler et al., 2004, Keunen et al., 2011). Therefore, it is important to understand how plants respond to heavy metal stress physiologically and molecularly, and to develop plants that can resist stress-induced ROS so that crop productivity can be maintained.

The roles of glutathione (*GSH*) and phytochelatin synthase (*PCs*) genes in reducing heavy metals stress and ROS scavenging have also been documented (Millar et al., 2003; Freeman et al., 2004; Hirata et al., 2005; Foyer and Noctor, 2005; Shao et al., 2008). In addition, the roles of antioxidants in scavenging ROS and reducing the oxidative stress caused by heavy metals have also been investigated (Hirschi et al., 2000, Tseng et al., 2007). Generally, a higher antioxidant content is found in anthocyanin-enriched plants, and these plants can survive abiotic and biotic stress conditions (Winkel-Shirley, 2002; Dixon et al., 2005; Dehghan et al., 2014; Agati et al., 2011). In addition, it has been suggested that improved abiotic stress tolerance when flavonoid accumulation is enhanced may be linked to increased ROS scavenging abilities (Fini et al., 2011; Nakabayashi et al., 2014). Transgenic plants overexpressing the transcription factor IbMYB1 and Del showed enhanced anthocyanin production and improved abiotic tolerances (Cheng et al., 2013; Naing et al., 2017). The results of previous studies suggested that to overcome abiotic stress conditions, it is important to produce anthocyanin-enriched plants that can provide antioxidants. A study by Lim et al. (2016) indicated that overexpression of the RsMYB1 transcription factor enhanced anthocyanin and antioxidant activity. Ai et al. (2017) also showed that overexpression of RsMYB1 enhanced anthocyanin accumulation in petunia. However, they did not investigate the role of RsMYB1 in abiotic stress tolerance. Therefore, the functional role of RsMYB1 in abiotic stress tolerance needs to be characterized.

This study used transgenic petunia that expressed RsMYB1. The plants were developed for a previous study by Ai et al. (2016). The role of RsMYB1 in petunia tolerance to heavy metal stress was investigated by examining several factors. These were plant height, root length, fresh weight, uptake of heavy metals, and stomata density. The transcript levels of RsMYB1, and the genes related to antioxidants and heavy metals resistance were also investigated.

## Materials and Methods

### Plants materials

The transgenic petunia expressing RsMYB1 was developed by Ai et al. (2017). However, the T_2_seeds were used as the material source in this study. Briefly, the seeds were soaked in 0.05% sodium hypochlorite solution (Yuhan Co., Ltd., Seoul, South Korea) containing 0.01% Tween 20 (Duchefa, Haarlem, The Netherlands) for 10 minutes and then rinsed with sterile distilled water at least three times. The sterilized seeds were sown in MS basal medium containing 3% sucrose and 0.8% agar. The cultures were incubated at 25 ± 2°C with a 16-h photoperiod and a light intensity of 50 μmol m^−2^ s^−1^ for 30 days.

### Heavy metal treatments

The 30-day-old transgenic seedlings that were red and of uniform size were selected for the heavy metal stress experiment. The 30-day-old non-transgenic seedlings were used as the wild type (WT) plants. The seedlings, including the WTs, were then stressed by continuous culturing in MS liquid medium containing serial concentrations of CuSO_4_, ZnSO_4_, K_2_Cr_2_O_7_ (25 µM, 50 µM, and 100 µM), or MnSO_4_ (100 µM, 250 µM, and 500 µM) for 10 days on a rotary shaker set to 50 rpm. The MS liquid medium (without heavy metals) was used as the control. The culture conditions were the same as described above. Each treatment contained 20 seedlings and there were three replicates.

### Effects of the different heavy metals

After treating the seedlings with 100 µM of CuSO_4_, ZnSO_4_, K_2_Cr_2_O_7_, or 500 µM MnSO_4_ for 10 days, fifteen plants from the non-treated (control condition), and the treated transgenic and WT petunias were randomly selected from each treatment. These were then used to measure plant height, root length, fresh weight, stomata density, heavy metal uptake, and gene expression. The measurements were taken three times, and the data represents the means of three replicates

### RNA extraction and gene expression analysis by quantitative real time-PCR (qRT-PCR)

The transcript levels of RsMYB1, two stress tolerant genes (*GSH* and *PCs*), and three antioxidant genes (*SOD, CAT*, and *POX*) in RsMYB1-overexpressing plants and WT plants under the stress and control conditions were investigated. Total RNA was isolated from 100 mg of leaf tissue per treatment using TRI Reagent TM Solution (Ambion, USA). Exactly 1 μg of total RNA and an oligo dT20 primer were used for reverse transcription (ReverTra Ace-á, Toyobo, Japan). Then, the transcript levels of the genes (*GSH, PCs, SOD, CAT, POX, RsMYB1*) and ACTIN were measured using a StepOnePlus Real-Time PCR system (Thermo Fisher Scientific, Waltham, USA) (Naing et al., 2017). The primers and PCR conditions for the detected genes are listed in Table 1. Three samples per line were used, and the analysis was repeated three times.

**Table 1.**
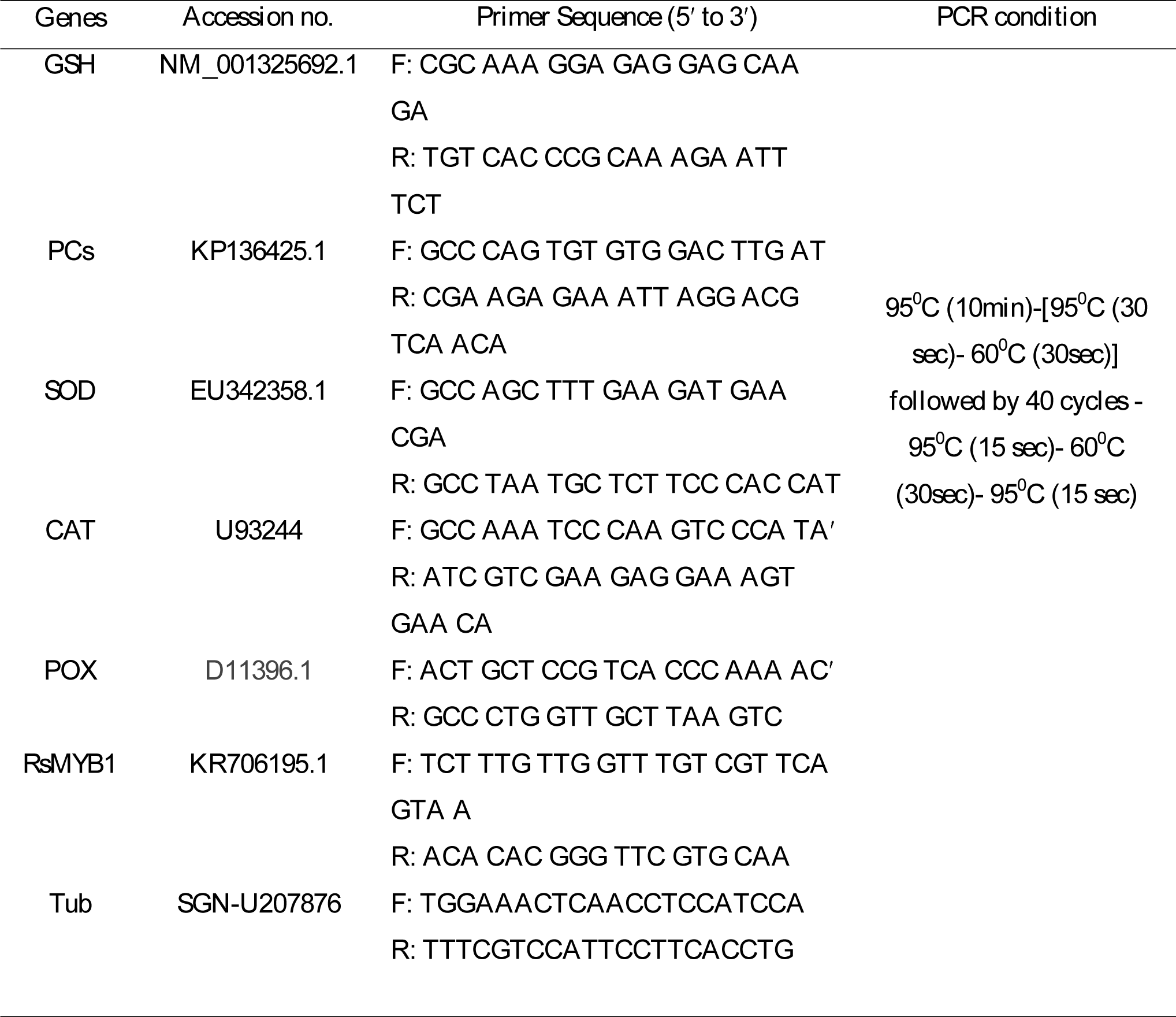
Primer sequences and PCR conditions used for qRT-PCR analysis in this experiment

### Uptake of the heavy metals

Exactly 1 g of dried leaf tissue per treatment was collected from RsMYB1-overexpressing plants and the WT plants to determine heavy metals (Cu, Zn, Mn, and Cr) uptake. The analysis was carried out as described by Cataldi et al. (2003). There were three samples per treatment and three replicates.

### Determination of stomata density by SEM

To determine whether the heavy metals treatments affected stomata density in RsMYB1-overexpressing plants and the WT plants, 5-mm-long leaf segments were excised with scalpel blades from the stress treatment and WT plants. The excised leaf segments were immediately fixed in formalin–acetic acid–alcohol and kept overnight according to the protocol used by Naing et al. (2015). The samples were then dehydrated for 10 min using serial ethanol concentrations (25, 50, 70, 85, and 100%). The dehydrated samples were dried to their critical point at room temperature, coated with gold-palladium on a Quick Cool Coater (Sanyu-Denshi, Japan), and the stomata density from each sample was examined using scanning electron microscopy (SEM; JEOL Ltd., Tokyo, Japan). Investigations were performed on three samples per treatment with three replicates.

### Statistical Analysis

Data were collected on the 30^th^ day after the starting date of the experiment and statistically analyzed using SPSS version 11.09 (IBM Corporation, Armonk, USA). The results are presented as means ± standard errors (SE). The Least Significant Different test (LSD) and t-test were used to separate the means, and the significance was set at P < 0.05.

## Results

### Assessment of plant growth parameters under heavy metal stress

Plant growth by the RsMYB1-overexpressing plants enriched with anthocyanin and the WT plants in response to the different heavy metals (CuSO_4_, ZnSO_4_, MnSO_4_, and K_2_Cr_2_O_7_) stresses, and the plants growing under normal growing conditions (without heavy metals) was evaluated 30 days after the treatments began. Under normal growing conditions, plant survival was high and they produced well-developed broad leaves and regular roots. Generally, there were no significant differences between the plants for the specific growth parameters, such as plant height, number of root and root length, and fresh weight. When they were treated with the 25 µM of CuSO_4_, ZnSO_4_, K_2_Cr_2_O_7_, or 100 µM MnSO_4_ for 10 days, their plant growth parameters were not significantly different from the plants grown under normal conditions (data not shown). However, these parameters started to decrease when the plants were treated with the 50 µM of CuSO_4_, ZnSO_4_, K_2_Cr_2_O_7_, or 250 µM MnSO_4_ for another 10 days (data not shown), and they significantly decreased when subjected to the 100 µM of CuSO_4_, ZnSO_4_, K_2_Cr_2_O_7_, or 500 µM MnSO_4_ treatment for a further 10 days (Figs. 1-4). In general, the RsMYB1-overexpressing plants were found to be more tolerant to heavy metal stress than the WT plants because the growth parameters evaluated in the RsMYB1-overexpressing plants were significantly higher than they were in the WT plants (Figs. 1–4). Some WT plants did not produce roots at all and their shoots showed signs of water deficiency and chlorophyll degradation (Fig. 5), but this was not observed in the RsMYB1-overexpressing plants. In addition, the degree of tolerance to the heavy metals by the plants varied depending on the heavy metal used. The CuSO_4_ and ZnSO_4_ compounds were found to be the most toxic to the plants, particularly the WT plants. Overall, the presence of high concentrations of the different heavy metals significantly inhibited the plant growth of the treated plants compared to the normal growing conditions. Furthermore, severe toxicity was clearly observed in the WT plants compared to the RsMYB1-overexpressing plants. Therefore, these results suggested that anthocyanin enriched RsMYB1-overexpressing plants had enhanced resistance to heavy metal stress.

**Fig. 1.**
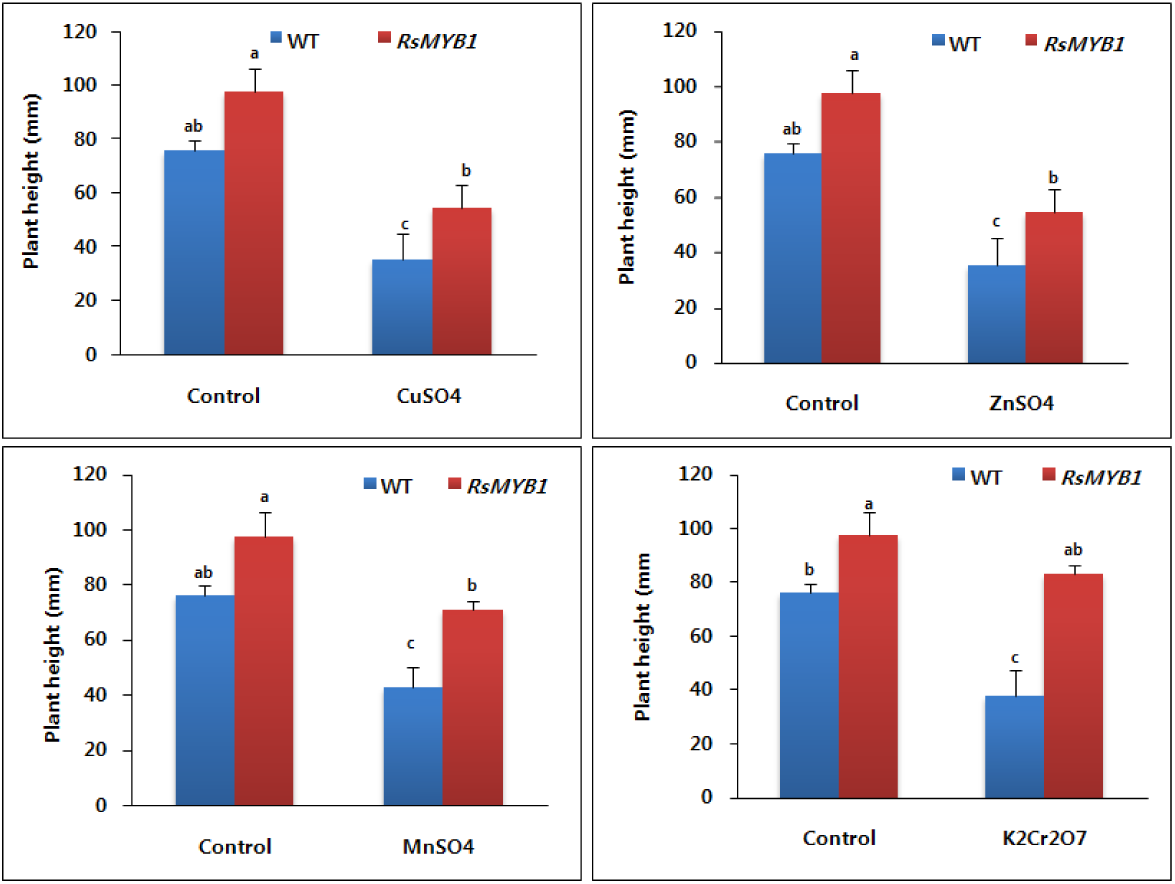
Comparisons of plant height of RsMYB1-overexpressing plant and WT under the different heavy metals stress. Data were taken on 30^th^ day after starting of the experiments. Error bars indicate the standard errors (SE) of average results

**Fig. 2.**
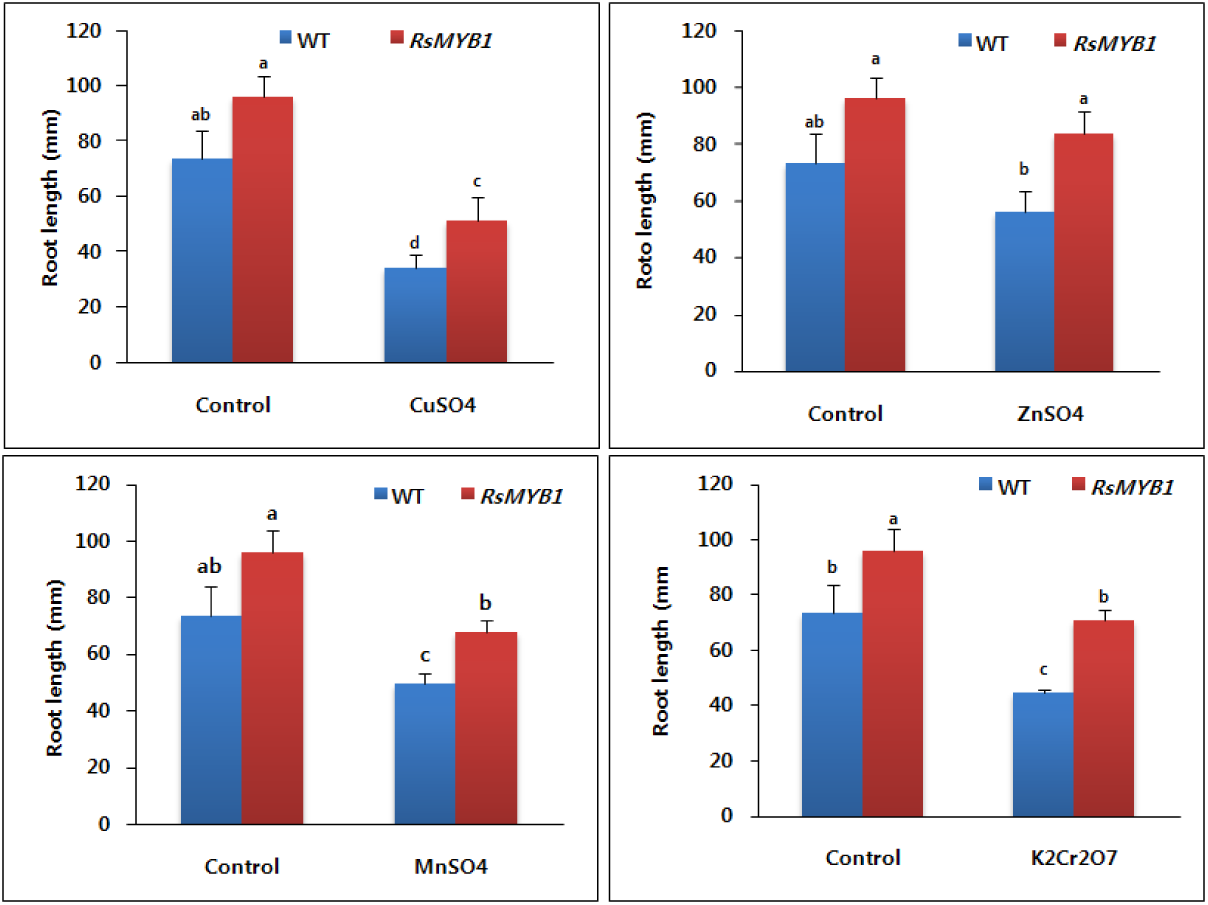
Comparisons of root length of RsMYB1-overexpressing plant and WT under the different heavy metals stress. Data were taken on 30^th^ day after starting of the experiments.. Error bars indicate the standard errors (SE) of average results

**Fig. 3.**
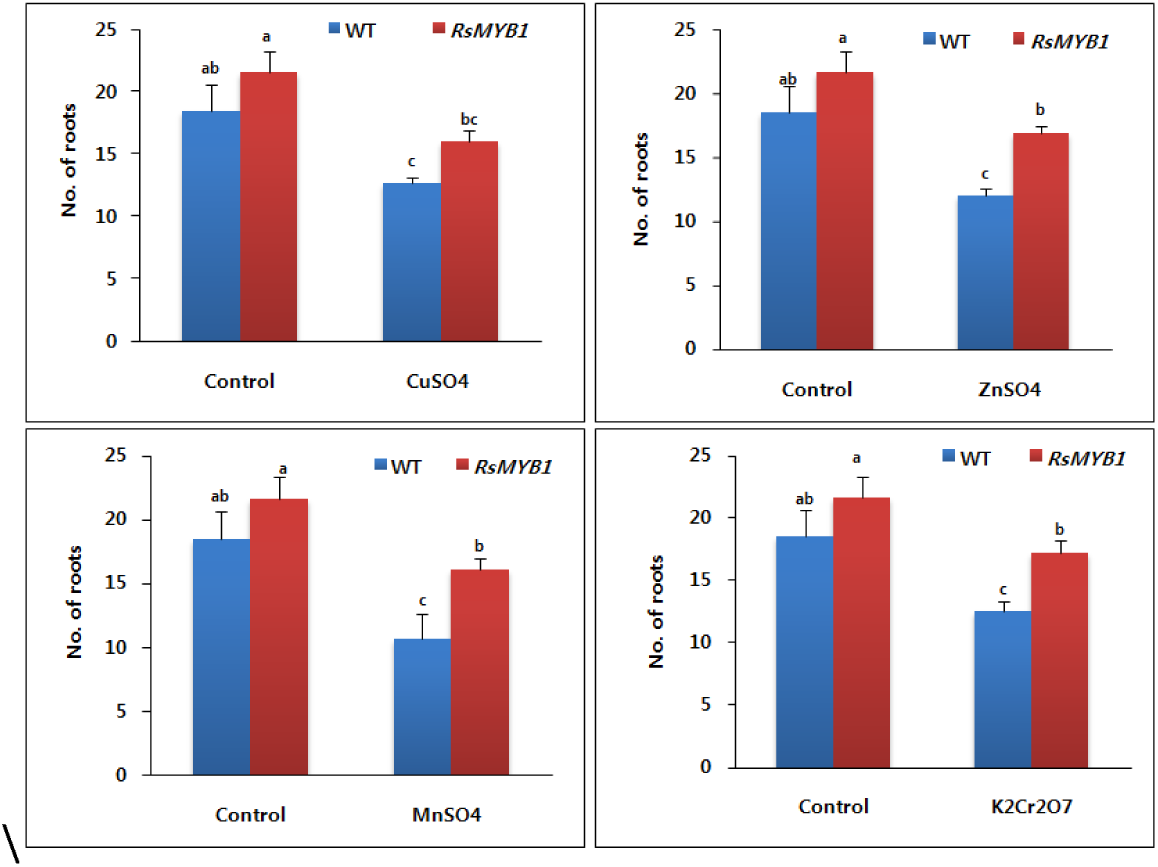
Comparisons of number of roots in RsMYB1-overexpressing plant and WT under the different heavy metals stress. Data were taken on 30^th^ day after starting of the experiments.. Error bars indicate the standard errors (SE) of average results

**Fig. 4.**
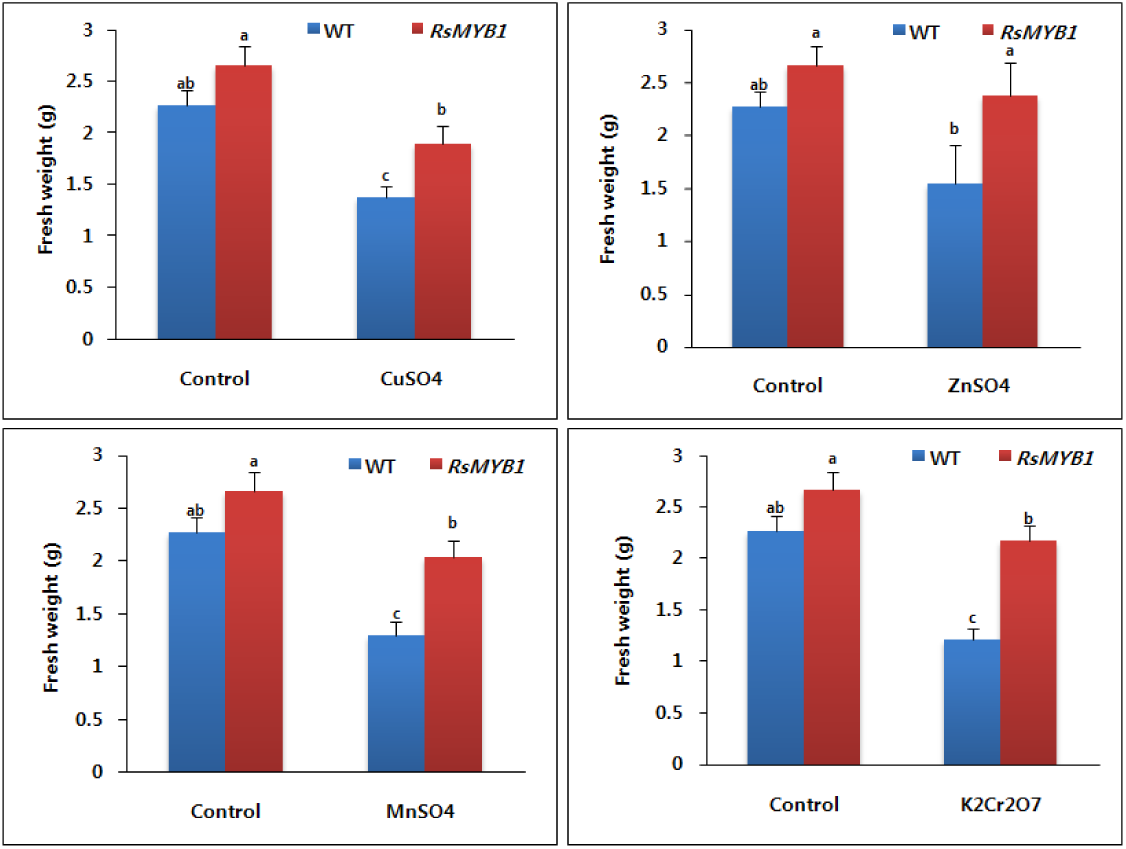
Comparisons of plant height of RsMYB1-overexpressing plant and WT under the different heavy metals stress. Data were taken on 30^th^ day after starting of the experiments.. Error bars indicate the standard errors (SE) of average results

**Fig. 5.**
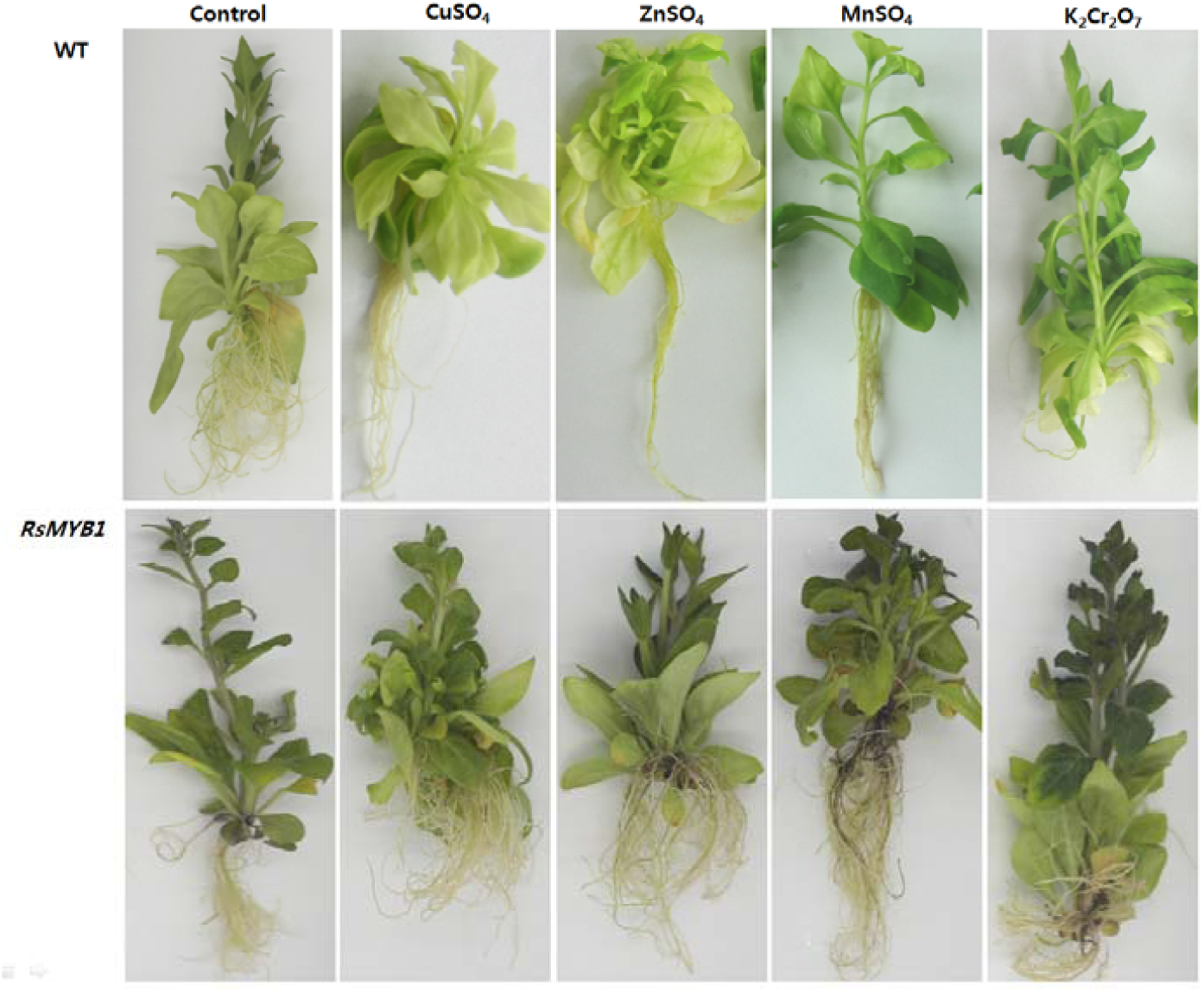
Comparisons of severity of toxicity in RsMYB1-overexpressing plant and WT caused by different heavy metals stress. Photo were taken on 30^th^ day after starting of the experiments.

### Reduction in stomata density under heavy metal stress

The stomata densities of the treated plants and the WT plants were investigated using a scanning electron microscope to determine whether the heavy metals affected the stomata density of the treated plants. Under normal growth conditions, a high stomata density was observed in both the RsMYB1-overexpressing plants and the WT plants. However, stomata density decreased when they were exposed to the heavy metals (Fig. 6). In addition, the extent of the reduction in stomata density varied depending on the type of heavy metal used.

**Fig. 6.**
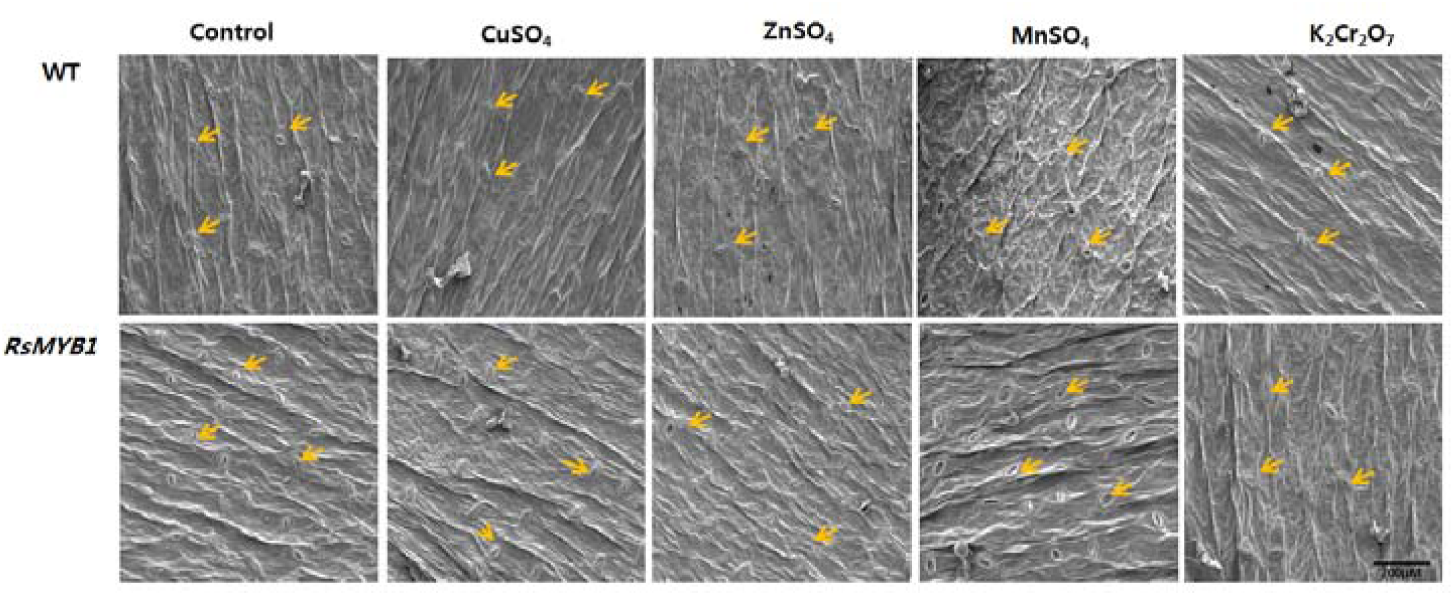
Comparisons of reduction of stomata densities in RsMYB1-overexpressing plant and WT caused by different heavy metals stress. Photo were taken on 30^th^ day after starting of the experiments

### Expression profiles of the antioxidant genes under heavy metal stress

The qRT-PCR method was used to clarify whether overexpression of RsMYB1 affects the expression levels of the antioxidant genes (*SOD, CAT*, and *POX*). The results showed that the stress treatments increased the transcript levels of the tested genes in the WT and RsMYB1-overexpressing plants compared to the normal growing conditions. However, the genes were more highly expressed in the RsMYB1-overexpressing plant than in the WT plants under all heavy metal stress conditions (Figs. 7–9). The expression levels of the genes were associated with the degree of tolerance to heavy metal stress because the RsMYB1 plants, which had higher expression levels of the antioxidant genes than the WT plants, were more tolerant to heavy metal stress than the WT plants. The results suggest that RsMYB1-overexpressing plants enriched with anthocyanin had higher antioxidant activities than the WT plants

**Fig. 7.**
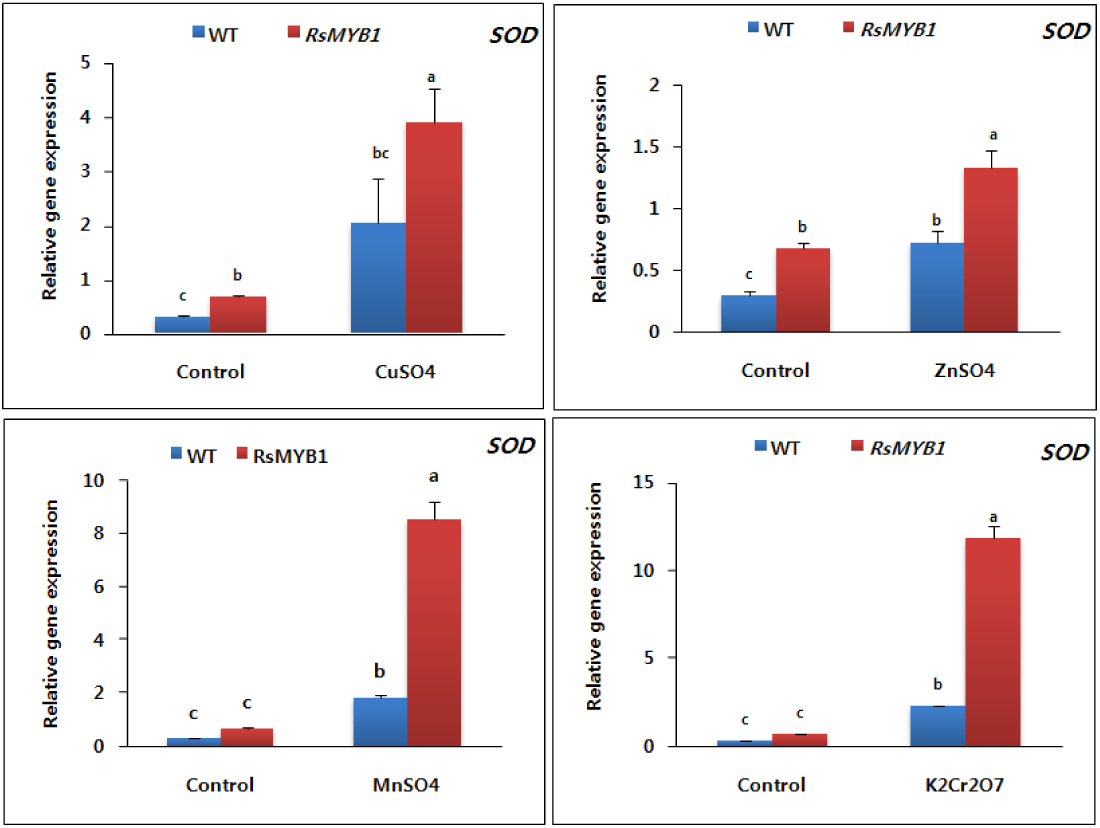
Expression analysis of the antioxidant-related gene (SOD) following the different heavy metals stress of WT and RsMYB1-overexpressing plants. Data were taken on 30^th^ day after starting of the experiments Error bars indicate the standard errors (SE) of average results

**Fig. 8.**
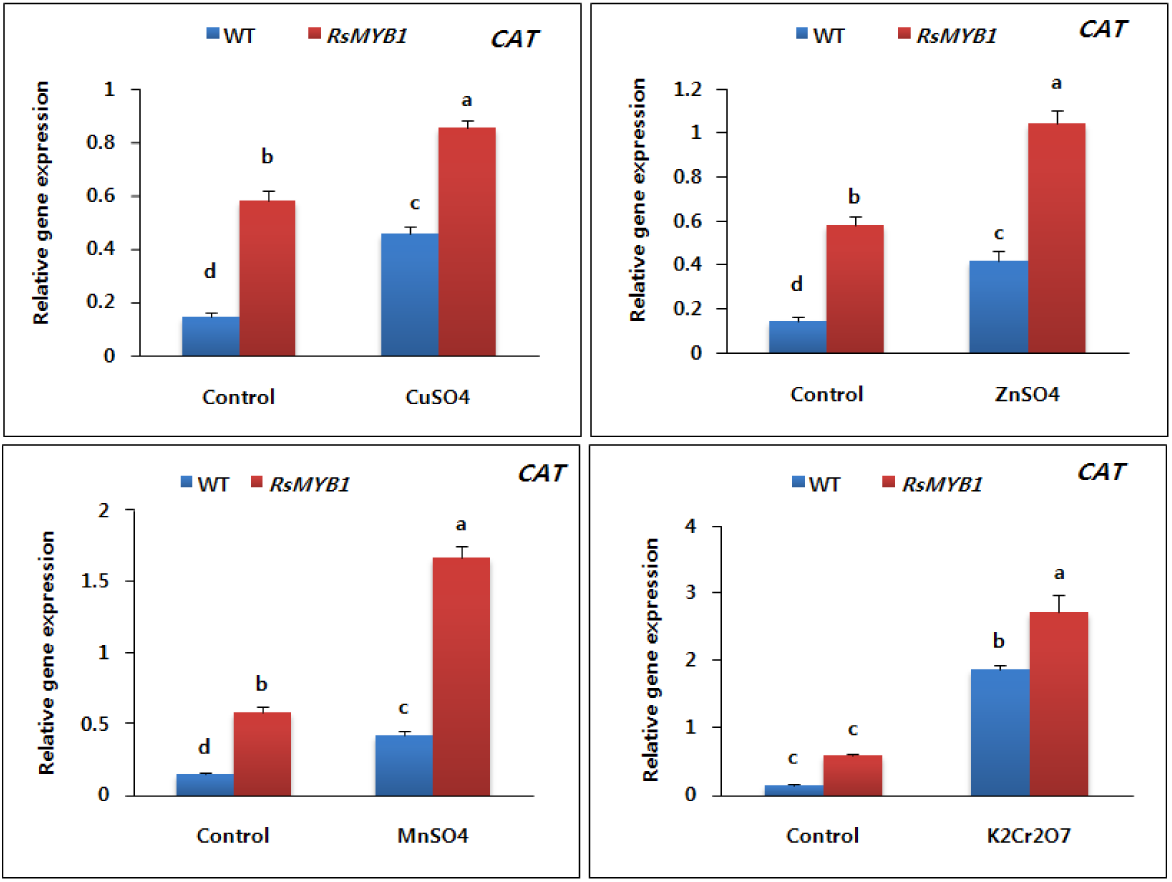
Expression analysis of the antioxidant-related gene (CAT) following the different heavy metals stress of WT and RsMYB1-overexpressing plants. Data were taken on 30^th^ day after starting of the experiments Error bars indicate the standard errors (SE) of average results

**Fig. 9.**
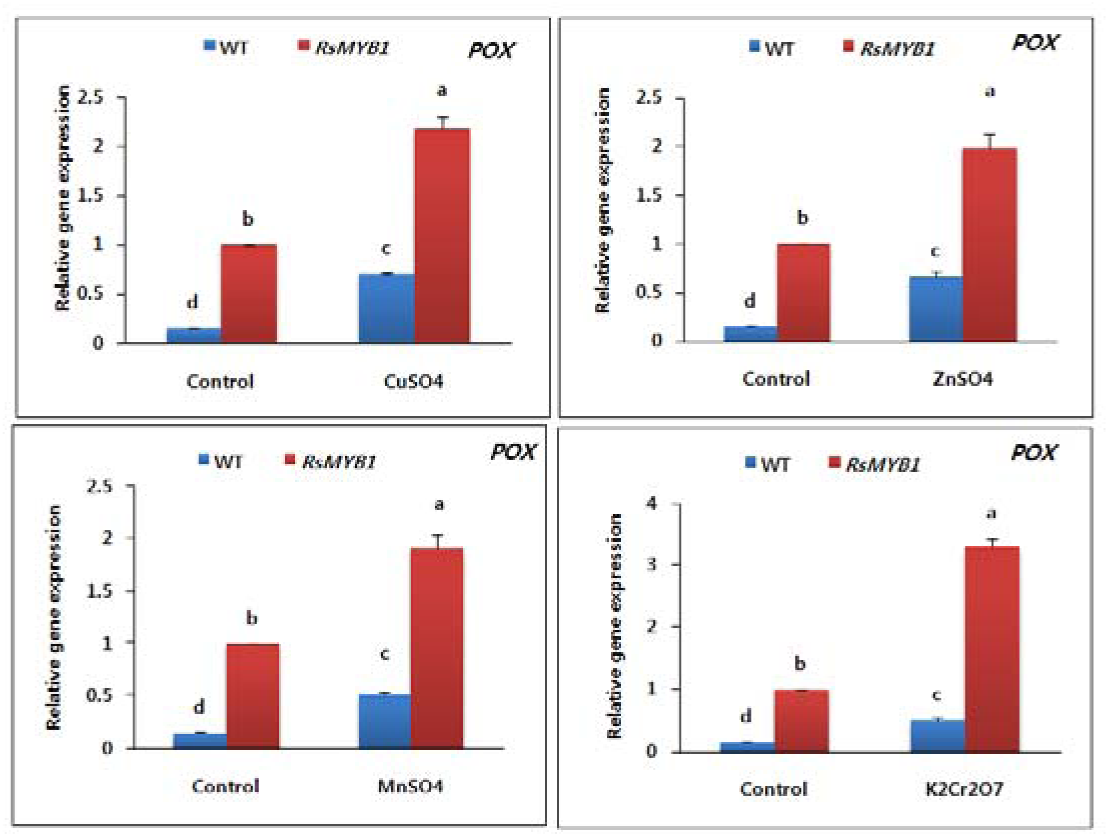
Expression analysis of the antioxidant-related gene (POX) following the different heavy metals stress of WT and RsMYB1-overexpressing plants. Data were taken on 30^th^ day after starting of the experiments Error bars indicate the standard errors (SE) of average results

### Expression profiles of the stress tolerant genes under heavy metal stress

The expression level responses of the stress tolerant genes (*GSH* and *PCs*), which are normally regulated by heavy metal stress, were analyzed by qRT-PCR. When the plants were subjected to heavy metal stress, their expression patterns were similar to the above-mentioned antioxidant gene patterns. The genes expressed in the treated and WT plants were transcriptionally low under normal growth conditions. When exposed to heavy metal stress, their expression levels were higher in the RsMYB1-overexpressing and WT plants. However, the stress-induced increase in expressions were higher in RsMYB1-overexpressing plants than in the WT plants (Figs. 10 and 11), which suggests that RsMYB1-overexpressing plants have a higher tolerance to heavy metal stress than the WT plants.

**Fig. 10.**
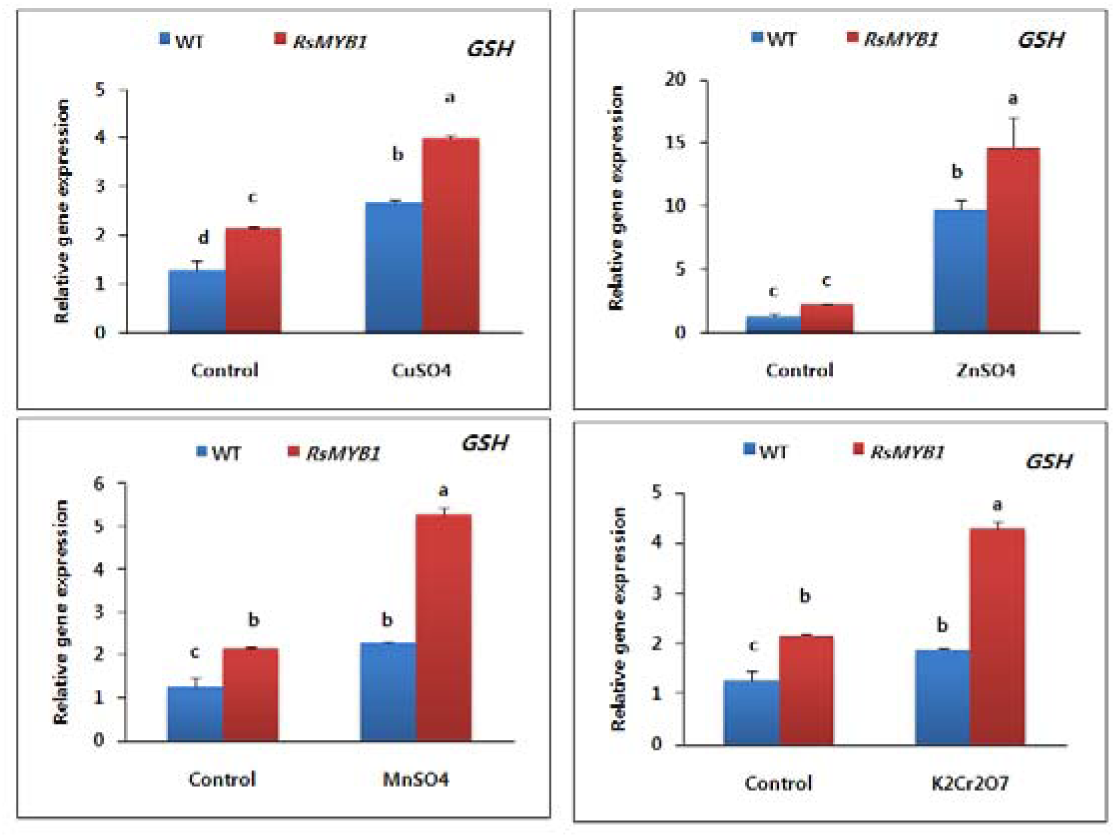
Expression analysis of the metal stress tolerant gene (GSH) following the different heavy metals stress of WT and RsMYB1-overexpressing plants. Data were taken on 30^th^ day after starting of the experiments Error bars indicate the standard errors (SE) of average results

**Fig. 11.**
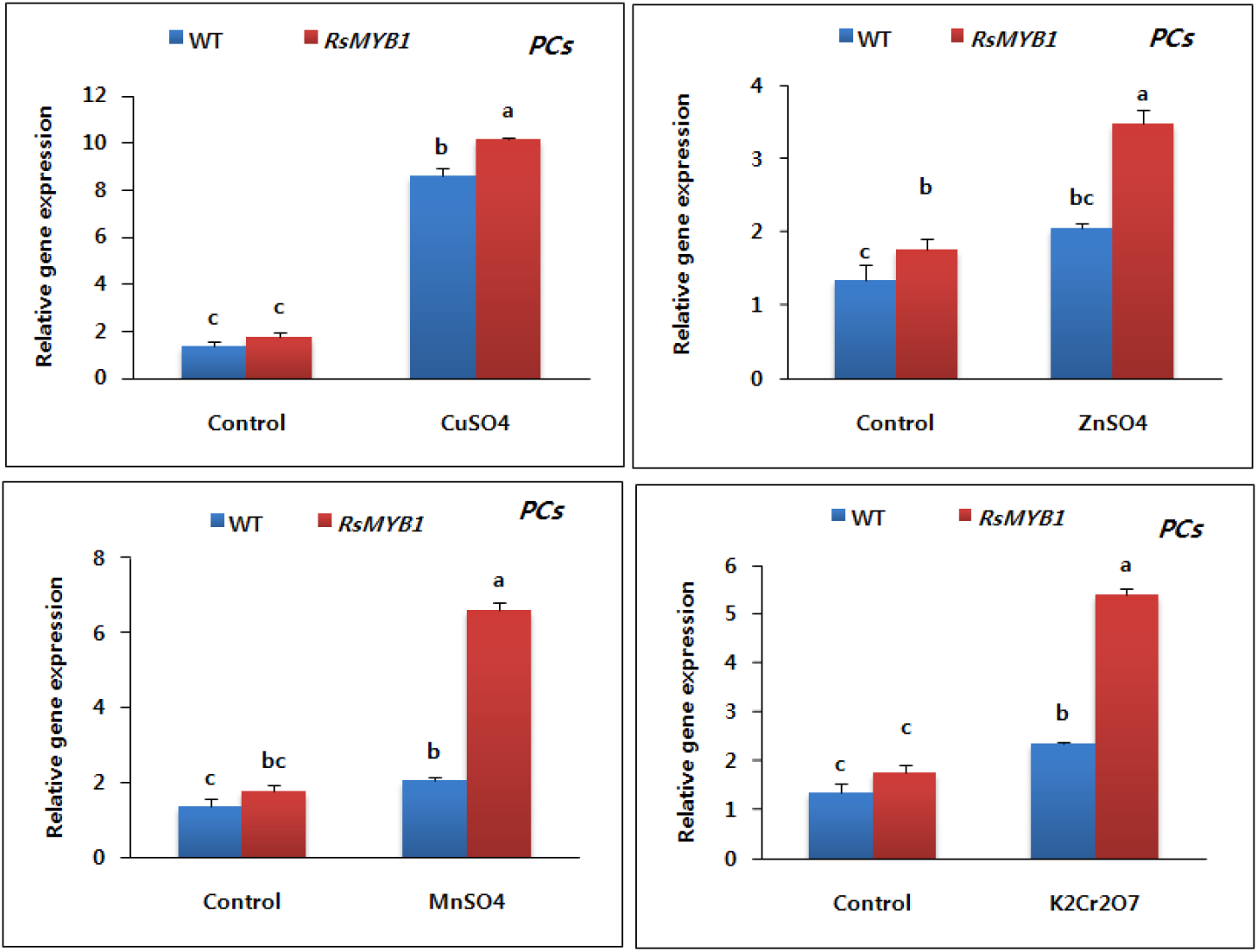
Expression analysis of the metal stress tolerant gene (PCs) following the different heavy metals stress of WT and RsMYB1-overexpressing plants. Data were taken on 30^th^ day after starting of the experiments Error bars indicate the standard errors (SE) of average results

### Accumulation of heavy metals in the RsMYB1-overexpressing and WT plants

Under normal growth conditions, the heavy metal levels in the overexpressing plants and the WT plants were quite low, and the two plant types were not significantly different. However, when they were exposed to the different heavy metals, the uptake of the metals by these plants significantly increased, but the total content of detectable metals per plant was significantly higher in the RsMYB1-overexpressing plants than in the WT plants (Fig. 12).

**Fig. 12.**
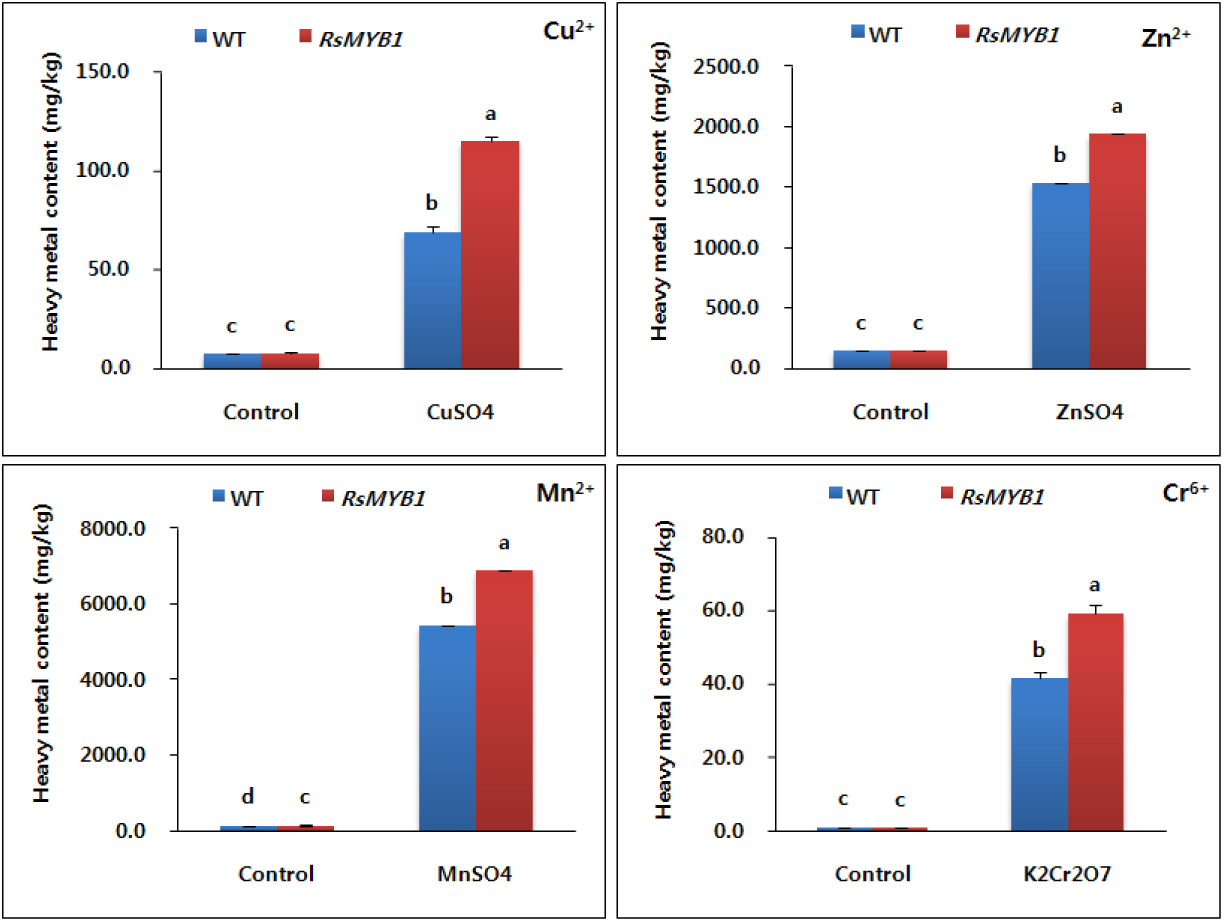
Accumulation of the heavy metal contents (Cu^2+^, Zn^2+^, Mn^2+^, Cr^6+^) in leaf tissue following heavy metal-stress treatment of WT and transgenic *RsMYB1*-overexpressing plants. Data were taken on the 30^th^ day after starting the experiments. Error bars indicate the standard errors (SE) of average results

### Elevation of RsMYB1 under heavy metal stress

As described above, RsMYB1-overexpressing plants were more tolerant to heavy metal stress than the WT plants because they could induce higher expression levels of stress- and antioxidant-related genes. Therefore, it was important to investigate whether the RsMYB1 expression level was elevated by heavy metal stress and whether its expression was associated with the regulation of the genes mentioned above. As expected, its expression level was elevated under the heavy metal stress conditions (Fig. 13), and this was linked to increased expressions of the stress- and antioxidant-related genes. This implies that RsMYB1 was involved in the regulation of the genes that are involved in abiotic stress defense. To clarify the role of RsMYB1 in abiotic stress tolerance, a phylogenetic tree was constructed based on the full-length amino acid sequences of thirty-five R2R3-MYB transcription factors isolated from different species that were found to be tolerant to different abiotic stresses (cold, drought, salt, and heavy metals). The resulting tree indicated that RsMYB1 was phylogenetically related to other MYB TFs and clustered with six TFs (IbMYB1, OsMYB4, GmMYB92, DwMYB2, OsMYB2, and TaMYB19) that confer different abiotic stress tolerances in various crops (Fig. 14). This suggested that RsMYB1 had the same functional role as the six TFs. According to the phylogenetic tree, RsMYB1 TF had strong sequence similarity with IbMYB1, which confers anthocyanin accumulation and salt-stress tolerance characteristics.

**Fig. 13.**
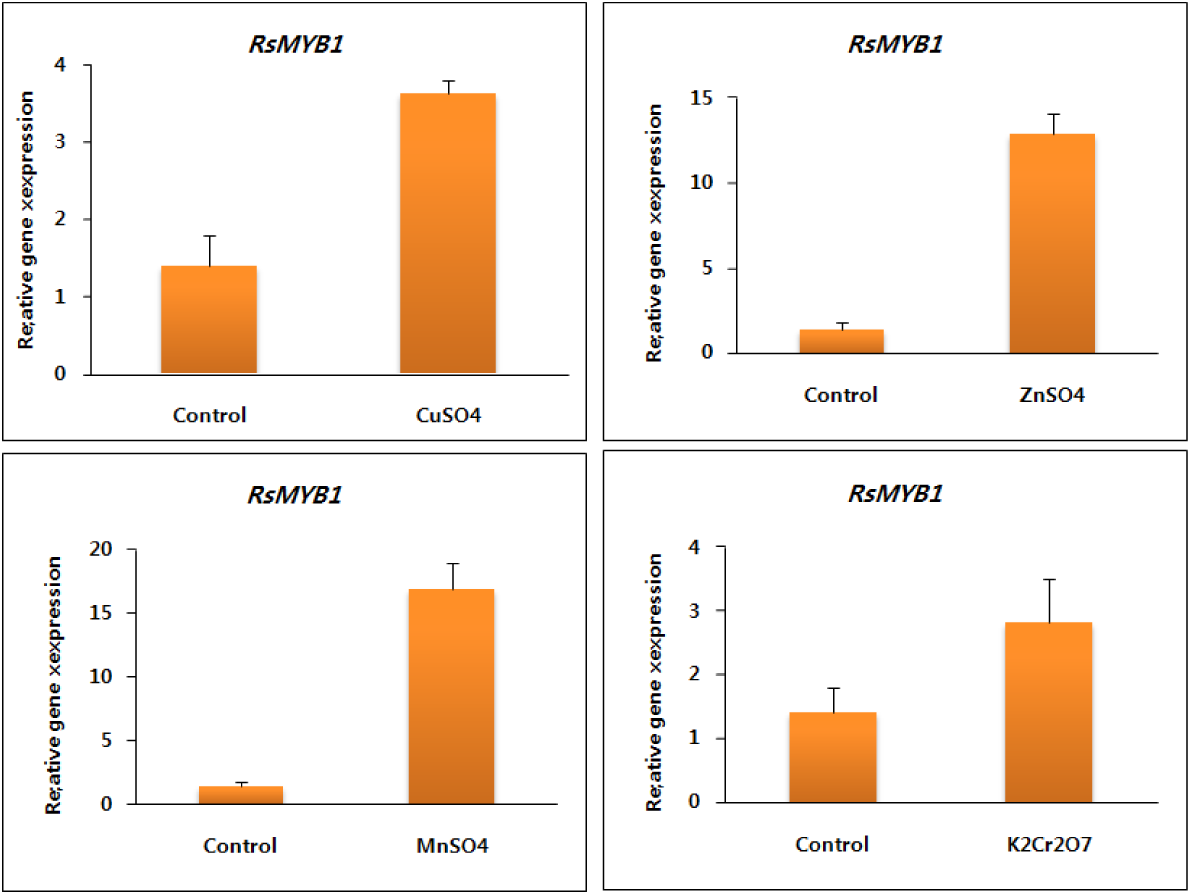
Expression analysis of RsMYB1 in RsMYB1-overexpressing plants following the different heavy metals stress. Data were taken on 30^th^ day after starting of the experiments Error bars indicate the standard errors (SE) of average results

**Fig. 14.**
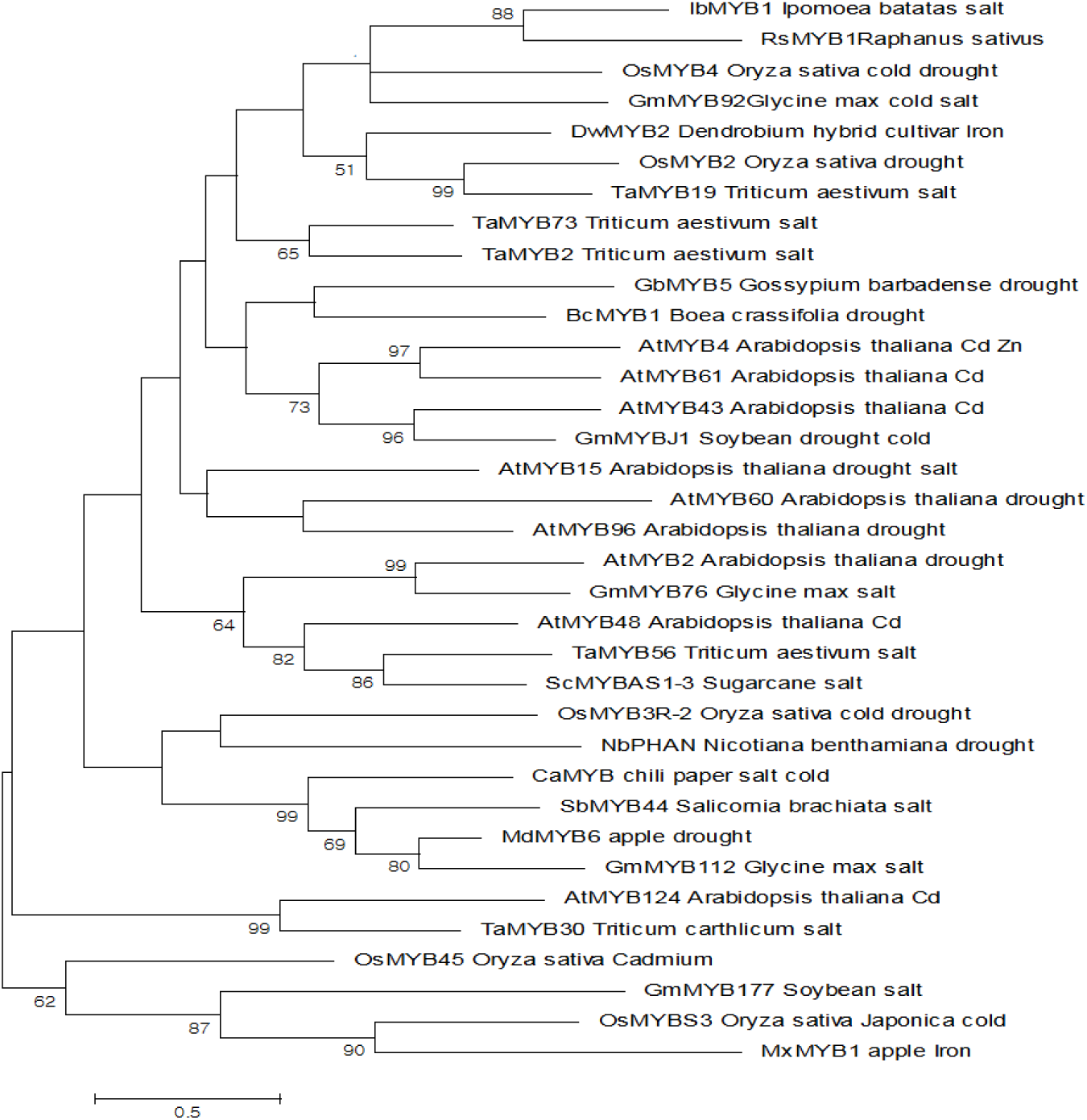
Phylogenetic relationships of the RsMYB1 transcription factor with other MYB TFs that confer different abiotic stress tolerances. The evolutionary history was inferred by using the Maximum Likelihood method based on the Poisson correction model [1]. The tree with the highest log likelihood (−4439.60) is shown. Initial tree(s) for the heuristic search were obtained automatically by applying Neighbor-Join and BioNJ algorithms to a matrix of pairwise distances estimated using a JTT model, and then selecting the topology with superior log likelihood value. The tree is drawn to scale, with branch lengths measured in the number of substitutions per site. The analysis involved 35 amino acid sequences. All positions containing gaps and missing data were eliminated. There were a total of 55 positions in the final dataset. Evolutionary analyses were conducted in MEGA7 [2].

## Discussion

Anthropogenic activities have led to a continuous increase in the agricultural soils contaminated by heavy metals. This is becoming a major concern worldwide (Rascio and Navari-Izzo, 2011; Villiers et al., 2011) because these contaminated soils negatively affect plant physiological processes through the generation of ROS, which results in lower crop yields (DalCorso et al., 2008; Hossain et al., 2009, 2010; Rascio and Navari-Izzo, 2011; Villiers et al., 2011). Therefore, there have been many studies on heavy metals toxicity and the defense mechanisms used by plants to scavenge ROS and detoxify heavy metals (Rascio and Navari-Izzo, 2011). Previous studies have shown that antioxidants are involved in the mechanism that scavenges the ROS generated by heavy metal stress (Mittler et al., 2004, Hirschi et al., 2000, Tseng et al., 2007). In addition, the roles of *GSH* and *PCs* in the detoxification of heavy metals and ROS scavenging have also been documented (Millar et al., 2003; Freeman et al., 2004; Hirata et al., 2005; Foyer and Noctor, 2005; Shao et al., 2008). The enhancement of antioxidant activities and stress tolerances in anthocyanin-enriched plants has been reported for various plant species (Winkel-Shirley, 2002; Dixon et al., 2005; Dehghan et al., 2014; Agati et al., 2011). Recently, overexpression of MYB TFs in various species has been shown to enhance anthocyanin accumulation and abiotic stress tolerance (Cheng et al., 2013; Meng et al., 2014; Qi et al., 2015; Yuan et al., 2015). In addition, Lim et al. (2016) also claimed that overexpression of RsMYB1 TF enhanced anthocyanin and antioxidant activity. In a previous study, overexpression of RsMYB1 in petunia enhanced anthocyanin accumulation (Ai et al., 2017). However, other functional roles of RsMYB1 in the abiotic stress tolerance process were not investigated. Therefore, it was important to characterize the functional role of RsMYB1 in heavy metal stress tolerance by using RsMYB1-overexpressing and WT plants to determine the plant growth parameters and the expression levels of the genes involved in the mechanism that underlies tolerance to heavy metal stress.

In this study, RsMYB1-overexpressing and WT plants survived well under normal growth conditions, and the growth parameters (plant height, root length, and fresh weight) were not generally significantly different between the over-expressing and WT plants. However, when they were exposed to the different heavy metals (CuSO_4_, Zn SO_4_, MnSO_4_, and K_2_Cr_2_SO_4_), there was a significant inhibition of the plant growth parameters for both plant types, especially with the WT plants. This was consistent with the results of previous studies, which reported that heavy metals were toxic to plants and caused injuries through the generation of ROS (Ebbs and Kochian, 1997; Lewis et al., 2001; Sharma et al., 2003; Scoccianti et al., 2006). In addition, the toxicity caused by CuSO_4_ and ZnSO_4_ was more severe than the other heavy metals, because they strongly inhibited plant growth and caused leaf chlorosis. The occurrence of such symptoms after exposure to CuSO_4_ and ZnSO_4_ in this study confirmed the results reported by Ebbs and Kochian (1997) and Lewis et al. (2001). The increased tolerance to heavy metals shown by RsMYB1-overexpressing plants compared to the WT plants is probably due to the increased anthocyanin levels in the RsMYB1-overexpressing plants because higher anthocyanin accumulation has been linked to a greater ROS scavenging ability (Fini et al., 2011; Nakabayashi et al., 2014) Theoretically, stomata play a critical role in photosynthesis because they are responsible for the uptake of carbon dioxide from the atmosphere and the release of oxygen as a waste product. In this study, inhibition of plant growth under heavy metal stress was associated with a reduction in stomata density. This suggested that the stomata density reduction caused by heavy metal stress decreased photosynthesis, which led to reduced plant growth. These results support the findings reported by Yilmaz et al. (2009) and Kambhampati et al. (2005), who also reported that heavy metals reduced stomata density.

The expression levels of the antioxidant genes (*SOD, CAT*, and *POX*) responsible for scavenging or neutralizing ROS were investigated to reveal the mechanisms which led to RsMYB1-overexpressing plants being more tolerant to heavy metal stress than WT plants. The expression levels of these genes were significantly higher under heavy metal stress than under normal growth conditions for both the overexpressing plants and the WT plants. This might be because the plants enhanced their expression levels to defend against ROS formation caused by the heavy metals. This supports the findings of previous studies (Bashri and Prasad, 2015; Singh et al., 2013). However, the greater inhibition of WT plant growth compared to the RsMYB1 overexpressing plants suggests that the antioxidant inducement in WT plants was significantly lower than in RsMYB1 overexpressing plants. Furthermore, gene inducement in the WT plants seems to have been insufficient to properly scavenge ROS. Under the same conditions, the higher expression of the genes in the RsMYB1-overexpressing plants than in the WT plants could be explained by the presence of anthocyanin, which is regulated by RsMYB1. High anthocyanin accumulation is linked to high antioxidant activities (Naing et al., 2017; Winkel-Shirley, 2002, Dixon et al., 2005, Dehghan et al., 2014; Agati et al., 2011). Therefore, this result supports the hypothesis that the higher stress tolerance in RsMYB1-overexpressing plants compared to WT plants depends on their antioxidant content.

The roles of other genes, such as *GSH* and *PCs*, which are involved in the antioxidant defense mechanism and heavy metal detoxification, were also investigated. When the overexpressing and WT plants were exposed to heavy metal stress, the *GSH* and *PCs* expression levels increased to defend against the heavy metals stress. Furthermore, the higher expression levels in the RsMYB1-overexpressing plants were positively associated with the degree of tolerance to heavy metals (RsMYB1 > WT). The high expressions of these two genes suggested that they played major roles in scavenging ROS, the detoxification of xenobiotics and heavy metals, and the maintenance of ionic homeostasis (Freeman et al., 2004, Foyer and Noctor, 2005; Hirata et al., 2005; Shao et al., 2008). In addition, the overexpression of these genes in other plants (*Brassica juncea, Arabidopsis, Populus canescens*, and *Nicotiana tabaccum*) also enhanced tolerance to heavy metal stress (Bittsánszkya et al., 2005; Cairns et al., 2006; Singla-Pareek et al., 2006; Gasic and Korban, 2007a,b). In this study, despite the enhanced expression of the *GSH* and *PC*s genes under heavy metal stress, there was still a reduction in plant growth when the plants were exposed to heavy metals, which indicated that the *GSH* and *PCs* levels were not high enough to completely defend against heavy metal stress, particularly in the WT plants. This strongly suggests that the heavy metal tolerance mechanism requires high GSH and PC levels.

More heavy metals accumulated in the RsMYB1-overexpressing plant shoots than in the WT plant shoots. This also suggests that the RsMYB1-overexpressing plants could be more tolerant to heavy metals than the WT plants, which led to the increased uptake of heavy metals. Another explanation for this is that the roots of the RsMYB1-overexpressing plants were less damaged by the metals than the WT roots because of the increased anthocyanin levels (higher antioxidant activities), and the higher expression levels of antioxidant (*SOD, CAT*, and *POX*) and metal-induced stress tolerant genes (*GSH* and *PCs*). This would have led to greater alleviation of the negative effects on root water uptake caused by metal stress. These results supported the findings of Zhu et al. (1999) and Wawrzyński et al. (2006), who reported that plants expressing the *GSH* and *PCs* genes showed enhanced Cd accumulation and tolerance. Bennett et al. (2003) also claimed that overexpression of *GSH* in mustard led the plant to accumulate 2.4-to 3-fold more Cr, Cu, and Pb than the wild-type plant.

In this study, the degree of stress tolerance was positively associated with the expression levels of antioxidant genes and stress tolerant genes. The expression levels of the antioxidant genes and stress tolerant genes were higher in RsMYB1-overexpressing plants than in WT plants under both growth conditions and particularly under heavy metal stress. Therefore, it was important to characterize the role of RsMYB1 and identify whether it was directly involved in the regulation of the genes, in addition to enhancing anthocyanin production. As expected, heavy metal stress elevated *RsMYB1* expression and this was linked to increased expressions of the stress- and antioxidant-related genes. Therefore, it is probable that RsMYB1 was not only responsible for enhancing anthocyanin production, but was also involved in the regulation of the genes participating in the mechanism that defends against abiotic stress. Recently, similar roles for MYB TFs have been reported (Cheng et al., 2013, Meng et al., 2014, Qi et al., 2015; Yuan et al., 2015). In addition, the role of RsMYB1 was further confirmed by results of the phylogenetic tree because RsMYB1 has strong phylogenetical relationships with six TFs (IbMYB1, OsMYB4, GmMYB92, DwMYB2, OsMYB2, and TaMYB19) that confer different abiotic stress tolerances in various crops.

## Conclusion

A previous study showed that RsMYB1 had a regulatory role during anthocyanin accumulation in petunia. However, its functional role in abiotic stress tolerance was not investigated. Therefore, this study characterized the functional involvement of RsMYB1 TF in tolerance to heavy metal stress using RsMYB1-overexpressing plants and WT plants. The results suggest that RsMYB1-overexpressing plants are more able to tolerate the heavy metal stress than WT plants because RsMYB1 induces higher expression levels of stress tolerant genes and antioxidant genes. In addition, it has a strong phylogenetic relationship with other MYB TFs that confer different abiotic stress tolerances. Therefore, these results suggest that RsMYB1 could be exploited as a dual function gene that will improve anthocyanin production and heavy metal stress tolerances in horticultural crops.

## Funding

This work was supported by the Korea Institute of Planning and Evaluation for Technology in Food, Agriculture, Forestry and Fisheries (IPET) through the Agri-Bio industry Technology Development Program, funded by the Ministry of Agriculture, Food and Rural Affairs (MAFRA) (grant #: 315002-5).

## References

Agati G, Biricolti S, Guidi L, Ferrini F, Fini A, Tattini M, et al. The 342 biosynthesis of flavonoids is enhanced similarly by UV radiation and root zone salinity in L. vulgare leaves. J Plant Physiol. 2011; 168:204–12.

Ai TN, Naing AH, Arun M, Jeon SM, Kim CK (2017) Expression of RsMYB1 in Petunia enhances anthocyanin production in vegetative and floral tissues. Scientia Horticulturae 214:58–65

Bashri G, Prasad SM (2015) Indole acetic acid modulates changes in growth, chlorophyll fluorescence and antioxidant potential of L. Trigonella foenum-graecum grown under cadmium stress. Acta Physiol Plant. 37:49

Bennett, L.E., Burkhead, J.L., Hale, K.L., Terry, N., Pilon, M., Pilon-Smits, E.A.H., 2003. Analysis of transgenic Indian mustard plants for phytoremediation of metal-contaminated mine tailings. Journal of Environmental Quality 32, 432–440.

Bittsánszkya, A., Kfmives, T., Gullner, G., Gyulai, G., Kiss, J., Heszky, L., Radimszky, L., Rennenberg, H., 2005. Ability of transgenic poplars with elevated glutathione content to tolerate zinc(2+) stress. Environmental Interaction 31, 251–254.

Cairns, N.G., Pasternak, M., Wachter, A., Cobbett, C.S., Meyer, A.J., 2006. Maturation of Arabidopsis seeds is dependent on glutathione biosynthesis within the embryo. Plant Physiology 141, 446–455

DalCorso G, S. Farinati, S. Maistri, and A. Furini, “How plants cope with cadmium: staking all on metabolism and gene expression,” Journal of Integrative Plant Biology, vol. 50, no. 10, pp. 1268–1280, 2008.

Davidson, J. F.; Schiestl, R. H. Mitochondrial respiratory electron carriers are involved in oxidative stress during heat stress in Saccharomyces cerevisiae. Mol. Cell. Biol. 21:8483–8489; 2001.

Dehghan S, Sadeghi M, Pöppel A, Fischer R, Lakes-Harlan R, Kavousi HR, Vilcinskas A, Rahnamaeian M, et al. Differential inductions of phenylalanine ammonia-lyase and chalcone synthase during wounding, salicylic acid treatment, and salinity stress in safflower, Carthamus tinctorius. Biosci Rep. 2014; 34:273–82

Dixon RA, Xie DY, Sharma SB. Proanthocyanidins: a final frontier in flavonoid research? New Phytol. 2005; 165:9–28

Ebbs, S.D., Kochian, L.V., 1997. Toxicity of zinc and copper to Brassica species: implications for phytoremediation. Journal of Environmental Quality 26, 776–781.

Fini A, Brunetti C, Di Ferdinando M, Ferrini F, Tattini M, et al. Stress-induced flavonoid biosynthesis and the antioxidant machinery of plants. Plant Signal Behav. 2011; 6:709–11

Fontes R L S and Cox F R 1998 Zinc toxicity in soybean grown at high iron concentration in nutrient solution. J. Plant Nutr. 21, 1723–1730.

Foyer, C.H., Noctor, G., 2005. Redox homeostasis and antioxidant signaling: a metabolic interface between stress perception and physiological responses. Plant Cell 17, 1866–1875.

Fozia A., Muhammad A. Z., Muhammad A., Zafar M. K. Effect of chromium on growth attributes in sunflower (Helianthus annuus L.) Journal of Environmental Sciences. 2008; 20(12):1475–1480.

Freeman, J.L., Persans, M.W., Nieman, K., Albrecht, C., Peer, W., Pickering, I.J., Salt, D.E., 2004. Increased glutathione biosynthesis plays a role in nickel tolerance in Thlaspi nickel hyperaccumulators. Plant Cell 16, 2176–2191

Gasic, K., Korban, S.S., 2007a. Expression of Arabidopsis phytochelatin synthase in Indian mustard (Brassica juncea) plants enhances tolerance for Cd and Zn. Planta 225, 1277–1285.

Gasic, K., Korban, S.S., 2007b. Transgenic Indian mustard (Brassica juncea) plants expressing an Arabidopsis phytochelatin synthase (AtPCS1) exhibit enhanced As and Cd tolerance. Plant Molecular Biology 64, 361–369

Hirata, K., Tsuji, N., Miyamoto, K., 2005. Biosynthetic regulation of phytochelatins, heavy metal-binding peptides. Journal of Bioscience and Bioengineering 100, 593–599.

Hirschi KD, Korenkov VD, Wilganowski NL & Wagner GJ (2000) Expression of Arabidopsis CAX2in tobacco. Altered metal accumulation and increased manganese tolerance. Plant Physiol. 124: 125–133

Hossain MA, M. Hasanuzzaman, and M. Fujita, “Up-regulation of antioxidant and glyoxalase systems by exogenous glycinebetaine and proline in mung bean confer tolerance to cadmium stress,” Physiology and Molecular Biology of Plants, vol. 16, no. 3, pp. 259–272, 2010.

Hossain MA, M. Z. Hossain, and M. Fujita, “Stress-induced changes of methylglyoxal level and glyoxalase I activity in pumpkin seedlings and cDNA cloning of glyoxalase I gene,” Australian Journal of Crop Science, vol. 3, no. 2, pp. 53–64, 2009.

Izaguirre-Mayoral M. L., Sinclair T. R. Soybean genotypic difference in growth, nutrient accumulation and ultrastructure in response to manganese and iron supply in solution culture. Annals of Botany. 2005; 96(1):149–158.

Kambhampati MS, Begonia GB, Begonia MFT, Bufford Y 2005. Morphological and physiological responses of morning glory (Ipomoea lacunosa L.) grown in a lead-and chelate-amended soil. International Journal of Environmental Research and Public Health 2(2): 299–303.

Keunen, E., Remans, T., Bohler, S., Vangronsveld, J., Cuypers, A., 2011. Metal-induced oxidative stress and plant mitochondria. Int. J. Mol. Sci. 12, 6894–6918.

Lewis, S., Donkin, M.E., Depledge, M.H., 2001. Hsp70 expression in Enteromorpha intestinalis (Chlorophyta) exposed to environmental stressors. Aquatic Toxicology 51, 277–291.

Meng X, Yin B, Feng HL, Zhang S, Liang XQ, Meng QW. Over-expression of R2R3-MYB gene leads to accumulation of anthocyanin and enhanced resistance to chilling and oxidative stress. Biol Plant. 2014; 58:121–30

Millar, A.H., Mittova, V., Kiddle, G., Heazlewood, J.L., Bartoli, C.G., Theodoulou, F.L., Foyer, C.H., 2003. Control of ascorbate synthesis by respiration and its implications for stress responses. Plant Physiology 133, 443–447

Mittler, R., Vanderauwera, S., Gollery, M., and Van Breusegem, F. (2004). Reactive oxygen gene network of plants. Trends Plant Sci. 9, 490–498

Naing AH, Park KI, Ai TN, Chung MY, Han JS, Kang YW, Lim KB, Kim CK (2017) Overexpression of snapdragon Delila (Del) gene in tobacco enhances anthocyanin accumulation and abiotic stress tolerance. BMC Plant Biol 17(1):65

Naing AH, Park KI, Chung MY, Lim KB, Kim CK (2015) Optimization of factors affecting efficient shoot regeneration in chrysanthemum cv. Shinma. Braz J Bot.39: 975–984

Prasad, A.S. (2012). Discovery of human zinc deficiency: 50 years later. J. Trace Elem. Med. Biol. 26, 66–69.

Qi L, Yang J, Yuan Y, Huang L, Chen P, et al. Overexpression of two R2R3-MYB genes from Scutellaria baicalensis induces phenylpropanoid accumulation and enhances oxidative stress resistance in transgenic tobacco. Plant Physiol Biochem. 2015;94:235–43. 31.

Rascio N and F. Navari-Izzo, “Heavy metal hyperaccumulating plants: how and why do they do it? And what makes them so interesting?” Plant Science, vol. 180, no. 2, pp. 169–181, 2011

Rezai K., Farboodnia T. Manganese toxicity effects on chlorophyll content and antioxidant enzymes in pea plant (Pisum sativum L. c.v qazvin) Agricultural Journal. 2008; 3(6):454–458.

Ricachenevsky, F.K., Menguer, P.K., Sperotto, R.A., Williams, L. E., and Fett, J.P. (2013). Roles of plant metal tolerance proteins (MTP) in metal storage and potential use in biofortification strategies. Front. Plant Sci 4:144

Scoccianti, V., Crinelli, R., Tirillini, B., Mancinelli, V., Speranza, A., 2006. Uptake and toxicity of Cr (Cr3+) in celery seedlings. Chemosphere 64, 1695–1703

Shanker, A.K., Cervantes, C., Loza-Tavera, H., Avudainayagam, S., 2005. Chromium toxicity in plants. Environment International 31, 739–753.

Shao, H.B., Chu, L.Y., Lu, Z.H., Kang, C.M., 2008. Primary antioxidant free radical scavenging and redox signaling pathways in higher plant cells. International Journal of Biological Science 4, 8–14.

Sharma, D.C., Sharma, C.P., Tripathi, R.D., 2003. Phytotoxic lesions of chromium in maize. Chemosphere 51, 63–68

Singh S, Parihar P, Singh R, Singh VP and Prasad SM (2015) Heavy Metal Tolerance in Plants: Role of Transcriptomics, Proteomics, Metabolomics, and Ionomics. Front. Plant Sci. 6:1143

Singh VP, Srivastava PK, Prasad SM (2013) Nitric oxide alleviates arsenic-induced toxic effects in ridged Luffa seedlings. Plant Physiol Biochem 71:155–163

Singla-Pareek, S.L., Yadav, S.K., Pareek, A., Reddy, M.K., Sopory, S.K., 2006. Transgenic tobacco overexpressing glyoxalase pathway enzymes grow and set viable seeds in zinc-spiked soils. Plant Physiology 140, 613–623

Tseng MJ, Liu CW, Yiu JC (2007) Enhanced tolerance to sulfur dioxide and salt stress of transgenic Chinese cabbage plants expressing both superoxide dismutase and catalase in chloroplasts. Plant Physiol Biochem 45:822–833

Villiers F, C. Ducruix, V. Hugouvieux et al., “Investigating the plant response to cadmium exposure by proteomic and metabolomic approaches,” Proteomics, vol. 11, no. 9, pp. 1650–1663, 2011.

Warne MS, Heemsbergen D, Stevens D, McLaughlin M, Cozens G, Whatmuff M, Broos K, Barry G, Bell M, Nash D, Pritchard D, Penney N (2008) Modeling the toxicity of copper and zinc salts to wheat in 14 soils. Environ Toxicol Chem 27:786–792

Wawrzyński, A., Kopera, E., Wawrzyńska, A., Kamińska, J., Bal, W., Sirko, A., 2006. Effects of simultaneous expression of heterologous genes involved in phytochelatin biosynthesis on thiol content and cadmium accumulation in tobacco plants. Journal of Experimental Botany 57, 2173–2182.

Williams L.E., and Pittman, J.K. (2010). “Dissecting pathways involved in manganese homeostasis and stress in higher plants,” in Cell Biology of Metals and Nutrients, Plant Cell Monographs, Vol.17, eds R. Helland R.R. Mendal (Berlin, Heidelberg: Springer-Verlag), 95–117.

Winkel-Shirley B. Biosynthesis of flavonoids 483 and effects of stress. Curr Opin Plant Biol. 2002;5:218–23.

Yang X, Feng Y., He, Z., and Stoffell, P. J (2005). Molecular mechanisms of heavy metal hyper accumulation and phytoremediation. J. Trace Elem. Med. Biol. 18, 339–353

Yilmaz K, Akinci IE, Akinci S (2009) Effect of lead accumulation on growth and mineral composition of egg plant seedlings (Solanum melongena). New Zealand J Crop Horticul Sci 37:189–199

Yuan Y, Qi L, Yang J, Wu C, Liu Y, Huang L, et al. A Scutellaria baicalensis R2R3-MYB gene, SbMYB8, regulates flavonoid biosynthesis and improves drought stress tolerance in transgenic tobacco. Plant Cell Tiss Organ Cult. 2015; 120:961–72

Zhu, Y.L., Pilon-Smits, E.A.H., Jouanin, L., Terry, N., 1999a. Overexpression of glutathione synthetase in Indian mustard enhances cadmium accumulation and tolerance. Plant Physiology 119, 73–79

Zou J., Wang M., Jiang W., Liu D. Chromium accumulation and its effects on other mineral elements in Amaranthus viridis L. Acta Biologica Cracoviensia Series Botanica. 2006; 48(1):7–12.

